# Temporal order of mutations influences cancer initiation dynamics

**DOI:** 10.1101/2021.01.10.426139

**Authors:** Hamid Teimouri, Anatoly B. Kolomeisky

**Affiliations:** Department of Chemistry, Rice University, Houston, Texas, United States; Center for Theoretical Biological Physics, Rice University, Houston, Texas, United States; Department of Chemical and Biomolecular Engineering, Rice University, Houston, Texas, United States; Department of Physics and Astronomy, Rice University, Houston, Texas, USA

## Abstract

Cancer is a set of genetic diseases that are driven by mutations. It was recently discovered that the temporal order of genetic mutations affects the cancer evolution and even the nature of the decease itself. The mechanistic origin of these observations, however, remain not well understood. Here we present a theoretical model for cancer initiation dynamics that allows us to quantify the impact of the temporal order of mutations. In our approach, the cancer initiation process is viewed as a set of stochastic transitions between discrete states defined by the different numbers of mutated cells. Using a first-passage analysis, probabilities and times before the cancer initiation are explicitly evaluated for two alternative sequences of two mutations. It is found that the probability of cancer initiation is determined only by the first mutation, while the dynamics is specified by both mutations. In addition, it is shown that the acquisition of a mutation with higher fitness before mutation with lower fitness increases the probability of the tumor formation but delays the cancer initiation. Theoretical results are explained using effective free-energy landscapes.

## Introduction

It is widely accepted that cancer tumors are caused by several genetic alterations, such as point mutations, unsuccessful DNA repairs and DNA methylations, in normal tissue cells [6,7,24,25]. These changes lead to abnormal growth of those cells that can no longer support the required functions of the tissue. The mechanistic origin of the increased competitiveness of cancer cells have been the focus of significant research efforts that identified the key features of tumors [6,11,24,25]. Traditionally, the cancer phenotype has been considered as a sum of complementary properties that each mutation drives separately [6]. These views, however, have been recently challenged by experimental observations suggesting that the order of mutations is also an important factor in cancer development [9,13,22]. But the microscopic picture behind the effect of the order of mutations in cancer remains largely unexplored.

There are now several examples of how different order of mutations lead to dramatically different outcomes in the cancer formation and development [3,8–10,14,18,22]. In the mouse model of adrenocortical tumors, it was observed that when the mutation in the oncogene Ras preceded the mutation in p53 gene this led to highly malignant tumors with strong metastatic properties. But the reverse order of mutations (p53 first and then Ras) produced only benign tumors [8]. Another striking example comes from the studies on chronic myeloproliferative neoplasms (MPNs), which are myeloid tumors that contain on average between 5 to 10 somatic mutations [22]. Two driver mutations, *TET2* and *JAK2*, were identified as the most common in these cancers. Blood cells analysis showed that patients with TET2-first mutations were a decade older than the patients with JAK2-first mutations. Moreover, it was observed that the order of mutation acquisition leads to different cancer types and probabilities of survival [22]. It was also shown in colorectal cancers using genome-sequencing studies that the type of the first mutation and the order in which mutations occur strongly affect the nature of deceases [13].

These observations stimulated discussions on how to explain why the mutation order alters cancer initiation and evolution [9,13]. Three possible mechanisms (not mutually exclusive) have been proposed [9]. One of them suggests that the first mutation might change the accessibility of specific genomic regions and this prevents the second mutation from activating or repressing this region. If the order of mutations is reversed the blocking does not happen. The second idea is that initially one mutation can induce rapid cell growth and differentiation, while starting from another mutation lead to much slower growth and less differentiation. The third mechanism argues that different starting mutations might lead to different cellular microenvironment, influencing this way the distinct disease evolution. These mechanisms, however, are qualitative, but the increasing amount of experimental data and the need to have new medical strategies to fight against the cancer require more quantitative understanding.

Theoretical models play an important role in uncovering the mechanisms of cancer initiation and progression [4,19,25]. They provide a critical quantitative link between the underlying microscopic processes and the appearance of malignant tumors. Due to these theoretical studies and a large number of other important experimental studies, we currently know that the cancer starts when cells accumulate certain number and types of genetic alterations. Significant efforts have been made for modeling the dynamics of mutation acquisition and how it is governed by relevant genetic parameters such as the rate of mutations, the size of the population of cells and the rate of mutations proliferation [1,5,12,16,19,20]. Recently we developed a new theoretical framework to evaluate the cancer initiation dynamics [23]. It is based on a discrete-state stochastic description of cancer initiation as a fixation of driver mutations in the tissues. It allows to explicitly evaluate the probabilities for the cancer to appear and the times before it happens, which are viewed as the fixation probabilities and the fixation times, respectively. This framework was applied for estimating the cancer initiation times from experimentally available data for different types of cancer.

In this paper, we present a theoretical model of cancer initiation that allows us to quantitatively explain the effect of different order of mutations. In our theoretical approach, the cancer appearance is viewed as a fixation of two different mutations. This so-called two-hit model assumes that the first type of mutations occupies the whole tissue before the second type of mutations starts and eventually fills the system. Using the method of first-passage processes, we explicitly evaluate the probability and the dynamics of cancer initiation. It is found that the order of mutations strongly influences both fixation probabilities and fixation times. We show that if the first mutation has a higher fitness than the second mutation this increases the overall probability of the cancer initiation, but, surprisingly, delays the formation of the tumor. These observations are quantified using effective free-energy landscapes for the underlying stochastic processes of cancer initiation.

### 1 Model

In our theoretical approach, the cancer initiation is viewed as an event that starts when two different mutations take over the healthy tissue as illustrated in Fig. 1. More specifically, we assume that originally in the well-mixed tissue compartment there are *N* normal stem cells and the total number of cells is always fixed. At some time, which we view as *t* = 0, one stem cell gets a first mutation with a probability *u*_1_. It is realistic to assume that the mutation rate is very low (*N*_*u*_1__ ≪ 1) [15,19], and the further transformations in the system are taking place only via stem cells divisions and removals to keep the total number of cells fixed. Normal cells divide with a speed *b*, while the cells with the mutation divide with a rate *r*_1_*b*. The parameter *r*_1_ is known as a fitness parameter, and it is equal to the ratio of the division rates for the mutated cell over the normal cell. It reflects the overall physiological influence of the mutation on cellular metabolism: if *r*_1_ < 1 the mutation is disadvantageous, *r*_1_ = 1 corresponds to a neutral effect, while for *r*_1_ > 1 the mutation is advantageous. The overall number of mutated cells in the fixed population of stem cells will be fluctuating, and there is a time when all cells would become mutated. This is known as a mutation fixation, and the system cannot have normal cells after that. At this moment, we assume that the second mutation will appear with a probability *u*_2_, which is also very small (*N*_*u*_2__ ≪ 1). The cells with one mutation divide with the rate *r*_1_*b*, while the cells with two mutations divide with the rate *r*_2_*b*. The fitness parameter *r*_2_ reflects the accumulative effects of both mutations present in the cell.

**Figure 1.**
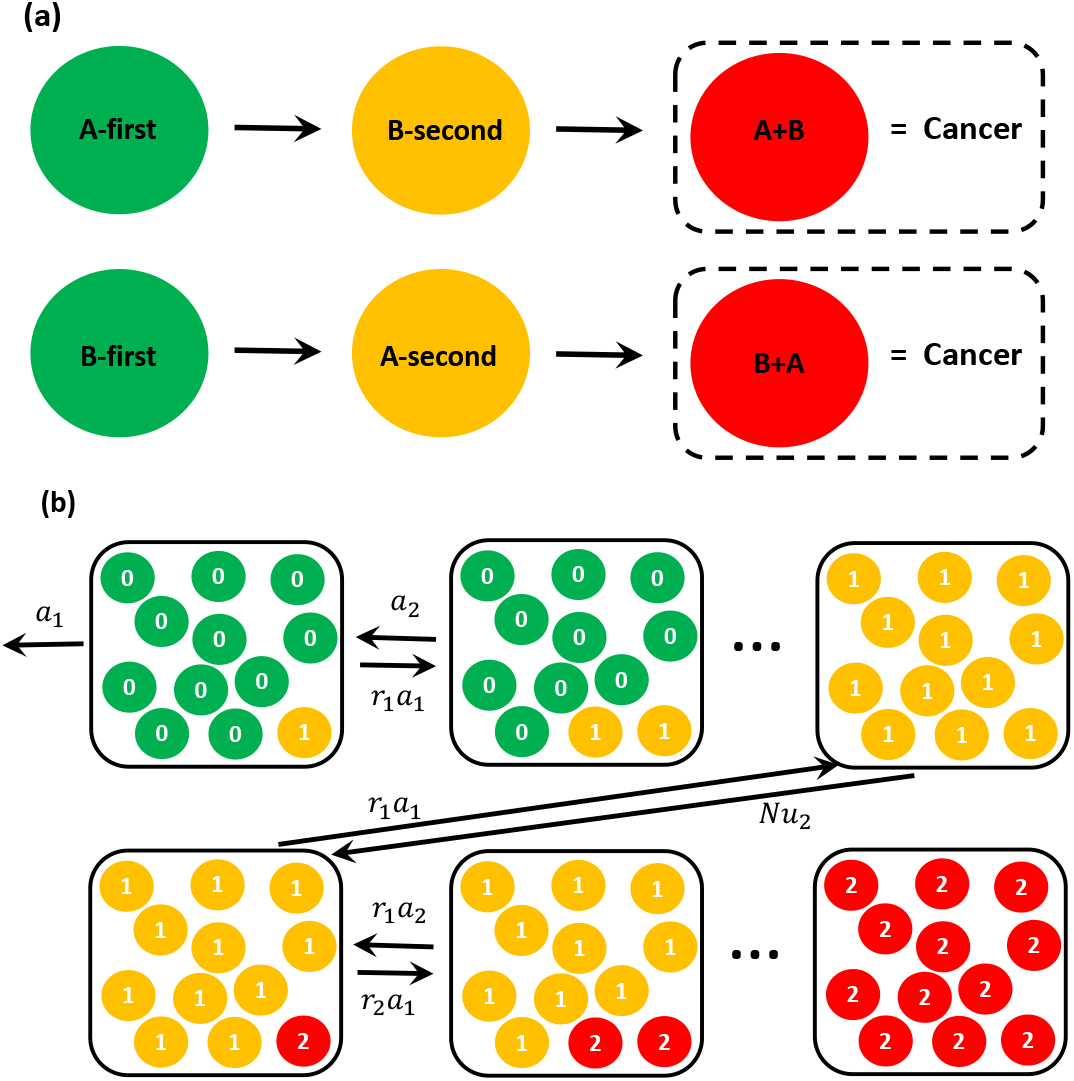
(a) A hypothetical scenario for the tissue to sequentially acquire two mutations, labeled *A* and *B*. The final outcome of two mutations is a cancer, but it might be different diseases depending on *AB* or *BA* sequences of mutations. (b) A schematic view of a two-hit mutation fixation process in the tissue compartment with *N* cells. Normal stem cells are green, while cells harboring single and double mutations are shown in yellow and red, respectively.

Introducing two mutations in the tissue compartment generates three types of cells (see Fig. 1b). The normal wild-type cells are labeled as type 0 (green circles), the cells with one mutation are labeled as type 1 (yellow circles), and the cells harboring two mutations are labeled as type 2 (red circles). At any time, the tissue might only have the cells of type 0 and 1, or only the cells of type 1 and 2 (Fig. 1b). The dynamic changes in the system can be viewed as stochastic transitions between 2*N* discrete states, as shown in Fig. 2, and these states are specified by the number of cells with one or two mutations. We define a state *n* (1 ≤ *n* ≤ *N*) as the state that has *n* cells with one mutation and *N* – *n* wild-type cells without mutations. If *N* < *n* ≤ 2*N* then the state *n* describes a situation with *n* – *N* cells with two mutations and 2*N* – *n* cells with one mutations: see Fig. 2. The state *n* = *N* corresponds to the fixation of the first mutation (all cells have one mutation), while the state *n* = 2*N* corresponds to the fixation of both mutations (all cells have two mutations). It can be shown that for the states on the first branch (1 ≤ *n* ≤ *N*), the forward transition rate from the state n to the state *n* + 1 is given by *r*_1_*a_n_* and the backward transition rate from the state *n* to the state *n* – 1 is equal to *a_n_* where, assuming that divisions and removals follow the Moran process [17], one obtains [23]

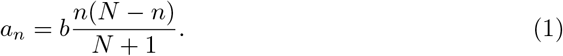

**Figure 2.**
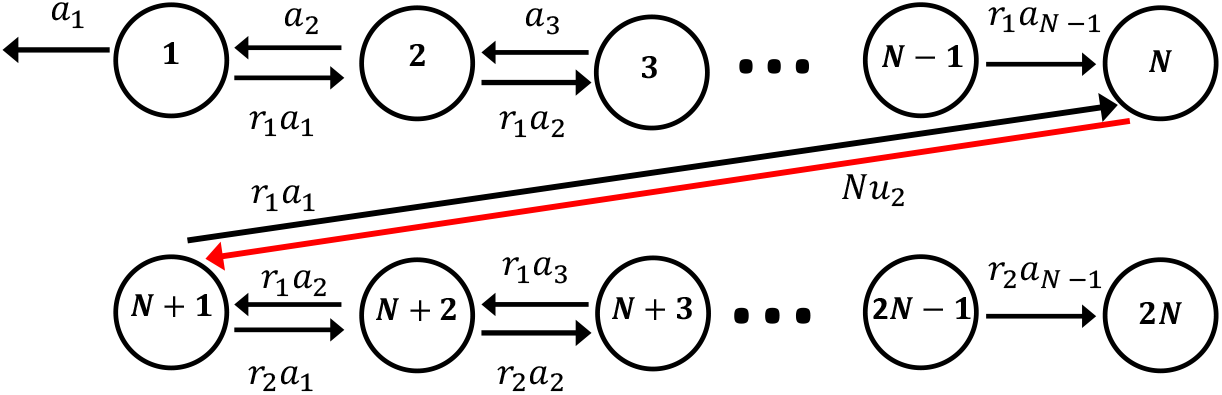
A discrete-state stochastic model for two sequential mutations in the tissue compartment. The state n corresponds to *n* cells with one mutations for 1 ≤ *n* ≤ *N*, and *n* – *N* cells with two mutations for *N* < *n* ≤ 2*N*.

For the states on the second branch (*N* < *n* ≤ 2*N*) it can be shown that the forward transition rate from the state n to the state *n* + 1 is given by *r*_2_*a_n–N_* and the reverse transition from the state *n* to the state *n* – 1 is given by *r*_1_*a_n–N_*: see Fig. 2. The transition rate from the state *N* to the state *N* + 1, which is given by the appearance of the second mutation, is equal to *N*_*u*_2__. Transitions between all neighboring states are reversible except for two situations. When the system goes from the state *N* – 1 to *N*, this corresponds to the elimination of all wild-type cells and it cannot be reversed. Similarly, the transition from the state 2*N* – 1 to 2*N* corresponds to the elimination of all cells with only one mutation. In addition, the single mutated cell can be eliminated from the system with the rate a_1_ from the state 1 (Fig. 2).

The dynamics of cancer initiation with two sequential mutations can now be explicitly analyzed using the method of first-passage processes [23]. This is because in our discrete-state stochastic model (Fig. 2) the cancer initiation corresponds to the events that start in the state 1 and reach the state 2*N* for the first time. We define a function *F_n_*(*t*) as the corresponding first-passage probability density function to start from any site *n* at *t* = 0 and to reach the final fixation state 2*N* at time *t*. The temporal evolution of these functions are governed by backward master equations,

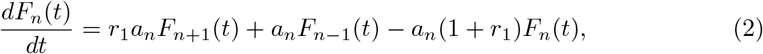

for 1 ≤ *n* < *N*, and

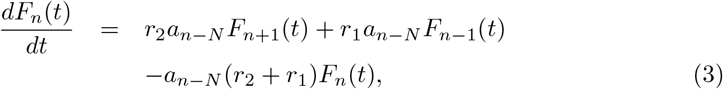

for *N* < *n* < 2*N*, while

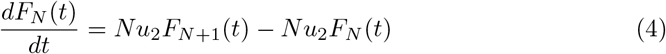

for *n* = *N*. In addition we have the boundary condition, *F*_2*N*_(*t*) = *δ*(*t*), which physically means that if the system starts in this state the cancer initiation process is immediately accomplished.

These equations can be solved explicitly (see the SI), and this allows us to obtain a comprehensive description of the cancer initiation dynamics. One can define a probability to reach the fixation state starting from the state *n*, 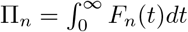, and the calculations presented in the SI show that

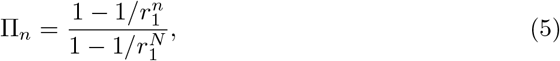

for 1 ≤ *n* < *N*, while Π_*n*_ = 1 for *N* ≤ *n* < 2*N*. This result can be easily understood by considering the properties of the discrete-state stochastic model in Fig. 2. If the system is found in one of the states 1 ≤ *n* < *N* then there is always a non-zero chance that the mutated cells will be eliminated from the system (exiting eventually to the left from the state 1 with the rate *a*_1_). However, because the transition from the state *N* – 1 to the state *N* is irreversible, for the states *N* ≤ *n* < 2*N* the system can never eliminate the mutated cells and with a probability one it will reach the fixation state. But there is another surprising observation from Eq. (5). It suggests that the probability of cancer initiation by two mutations (starting from the state *n* =1) depends only on the properties of the first mutation. This is again the consequence of the irreversibility in the transition from the first branch and transient fixation of the first mutation before the system can proceed further to the final fixation of both mutations. Since the probability of cancer initiation is directly related to the cancer lifetime risk [19,21], this clearly indicates that the order of mutations does matter. The results for the fixation probability also simplify in several limiting cases. When *r*_1_ → 1, we obtain Π_*n*_ = *n/N*, while for the tissues with very large number of cells and advantageous first mutation (*N* » 1 and *r*_1_ > 1) we have 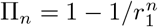.

Another important characteristics of the cancer initiation process is the cancer initiation time. It is defined as a time for the tissue to reach the two-mutations fixation for the first time. In our theoretical framework, it also corresponds to the mean first-passage time to reach the state 2*N* starting from the state *n*. It is generally defined as 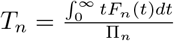. As shown explicitly in the SI, the cancer initiation time starting from the state *n* = 1 can be written as

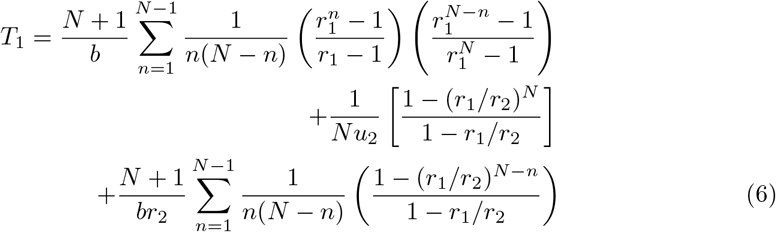

There are three contributions to the average fixation time of two mutations in Eq. (6), i.e., *T*_1_ = *T*_11_ + *T_trans_* + *T*_12_. The first term *T*_11_ describes the time for the system to reach the state *N*, which corresponds to the fixation of the first mutation. The second term *T*_trans_ describes the effective rate of acquiring the second mutation in cells that are fully fixed by the first mutation. It can be shown that

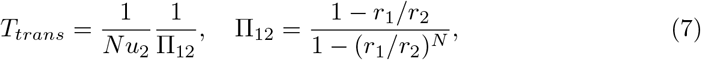

where Π_12_ is the fixation probability of the second mutation starting from the state *N* + 1. Thus, this contribution to the overall fixation time reflects the possibility of reverse transitions from the state *N* + 1 back to the state *N*. The third term, *T*_12_ describes the time to reach the final fixation starting from the state *N* + 1.

Again, it is interesting to consider limiting cases. For *r*_1_ = *r*_2_ = 1, i.e., for two successive neutral mutations and *N* → ∞, we obtain,

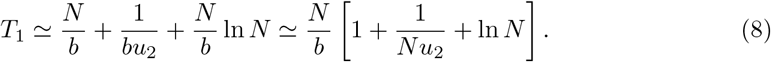

In this limit, the system spends most of the time on the second branch of discrete states. Also, for large *N* Eq. (6) can be simplified into (see the SI):

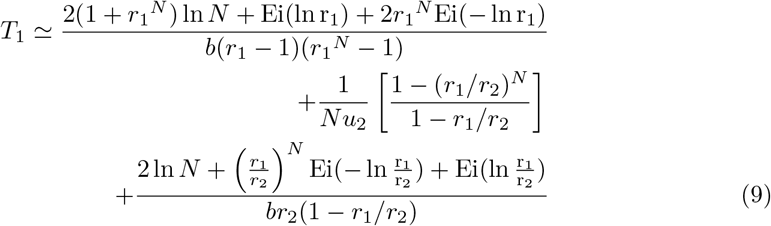

where Ei(*x*) is the exponential integral defined as 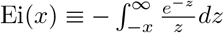, and *γ* is the Euler Mascheroni constant. For *r*_1_ < *r*_2_, terms multiplied by (*r*_1_/*r*_2_)^*N*^ vanish and consequently the fixation time is further simplified,

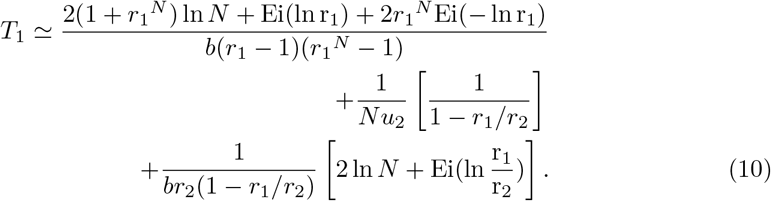

Now we can analyze the effect of the mutations order by considering two specific mutations *A* and *B* with the fitness parameters given by *r_A_* and *r_B_*, respectively. There are two different mutation sequences in this system, labeled as *AB* and *BA*. For the events of type *AB*, the mutation *A* is the first one, while the mutation *B* is the second one. Then we have *r*_1_ = *r_A_* while *r*_2_ = *εr_A_r_B_*. Here, the combined fitness parameter *r*_2_ reflects the presence of both mutations and the parameter *ε* describes the cooperativity between the mutations. When *ε* > 1 the presence of both mutations is more advantageous and this is a positive cooperativity, while for *ε* < 1 we have a negative cooperativity when the mutations counterbalance each other. To simplify our calculations, we assume *ε* = 1, which corresponds to a neutral cooperation, i.e., when the effect of both mutations is independent of each other. But the cooperativity effects can be easily taken into account in our theoretical approach. The situation is opposite for events of type *BA*. Here we have *r*_1_ = *r_B_* and *r*_2_ = *r_A_r_B_*. These two mutational sequences have different probabilities of cancer initiation defined as Π_*AB*_ and Π_*BA*_, respectively. To quantify the difference we define a ratio,

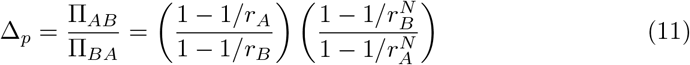

Fig. 3a shows the difference in the cancer initiation probabilities for different mutations sequences as a function of the relative fitness parameters. One can see that, when the first mutation is more advantageous (*r_A_*/*r_B_* > 1), the *AB* fixation probability is higher than the *BA* fixation probability. The effect is stronger when the fitness parameters are closer to being equal to one. This observation can be easily explained by noting again that the overall fixation probability depends only on the nature of the first mutation. Thus, if the first mutation has a larger fitness parameter for one sequence it will lead to a higher probability of the cancer initiation. Similarly, we can explicitly calculate the fixation times *T_AB_* and *T_BA_* for *AB* and *BA* mutation sequences, respectively. To quantify the difference, the following function is defined,

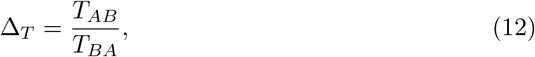

and the results are presented in Fig. 3b. It shows that there is a different cancer initiation dynamics depending on the mutations order, and the effect is stronger when fitness parameters are close to unity. However, the unexpected result is that if the first mutation is more advantageous (*r_A_*/*r_B_* > 1) it takes longer to initiate the cancer. This contrasts with the higher probability of the overall fixation for this situation, i.e., something that is more probable takes longer to achieve, which is opposite to naive expectations.

**Figure 3.**
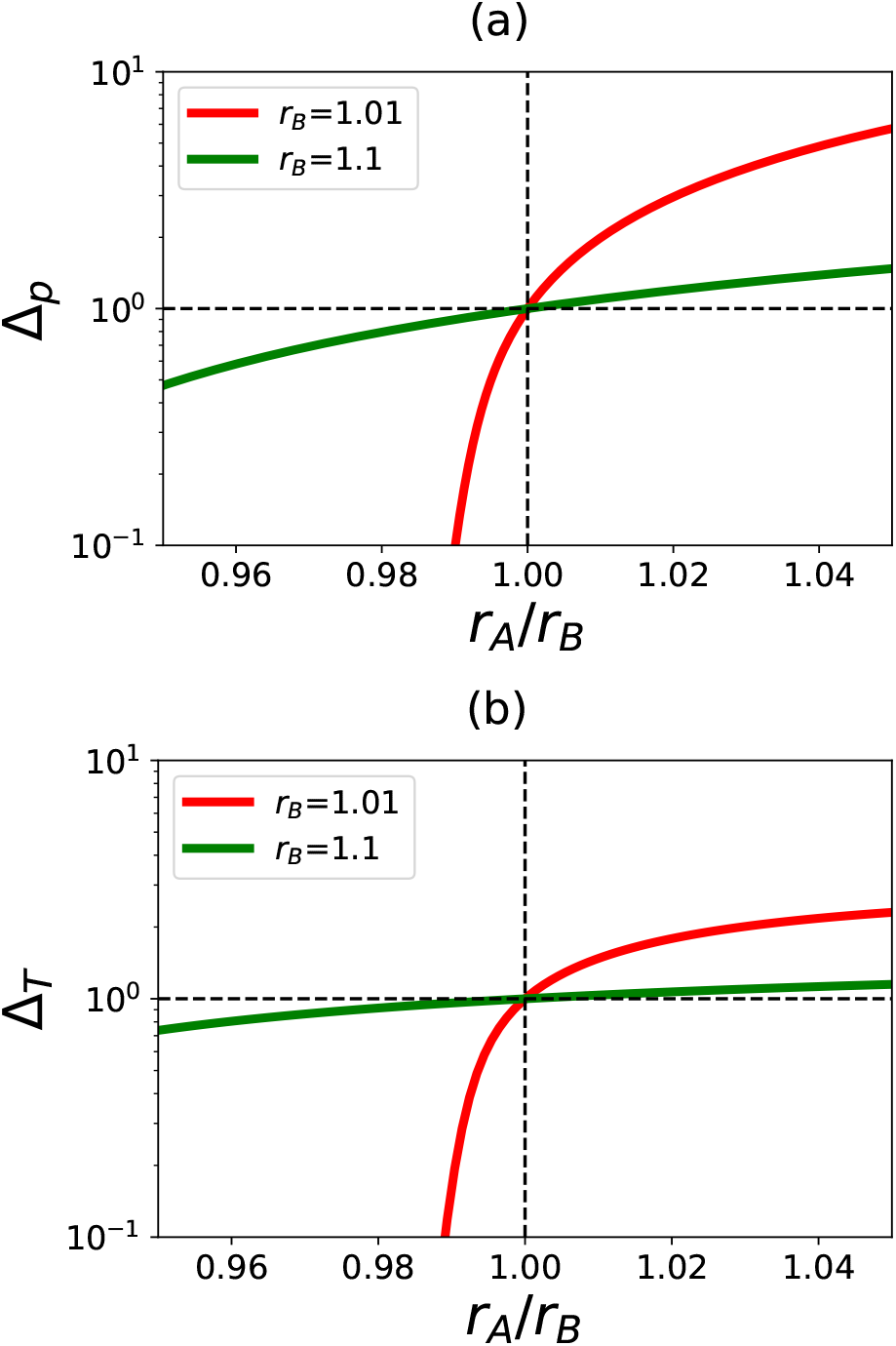
a) The ratio of fixation probabilities Π_*AB*_/Π_*BA*_ for two alternative sequences of mutations. b) The ratio of fixation times *T_AB_/T_BA_* for two alternative sequences of mutations. For calculations, *N* = 1000 is used.

To understand the microscopic origin of the surprising observations of anti-correlations between the probability and the time before the cancer initiation, we plot in Fig. 4 the ratio of the first and third terms in the fixation times (*T*_11_/*T*_12_) for different fitness parameters. These times correspond to the system being found on the first branch when only the wild-type cells and the cells with one mutation are present (*T*_11_, 1 ≤ *n* < *N*); and for being on the second branch when only the cells with one or two mutations are present (*T*_12_, *N* +1 ≤ *n* < 2*N*). For convenience, let us consider the *AB* mutation sequence. When the first mutation has a higher fitness parameter than the second mutation (Fig. 4, upper left corner), it is found that *T*_11_/*T*_12_ < 1, i.e., the system spends most of the time on the second branch of discrete states. But if the second mutation is more advantageous (Fig. 4, lower right corner) the situation is reversed, and the system spends most of the time on the first branch of the discrete states. Based on these observations, we can construct an effective “free-energy” landscape for the cancer initiation dynamics driven by stochastic transitions. One can associate longer times with higher barriers and shorter times with smaller barriers. This leads to a schematic picture shown in Fig. 5 for alternative sequences of mutations. The first barrier corresponds to the states on the first branch, and the second barrier corresponds to the states on the second branch. The deep region between two barriers reflects the irreversible transition between two branches of the states. Now we can understand better the difference in cancer initiation for two sequences of mutations. The probability of cancer initiation is determined only by the first barrier, and then it is clear that if the first mutation is more advantageous (*r_A_* > *r_B_*, left panel on Fig. 5) the cancer lifetime risk is higher. But in the opposite case (*r_B_* > *r_A_*, right panel in Fig. 5), the cancer lifetime risk is lower. At the same, the cancer initiation dynamics depends on both barriers, although the highest barrier dominates the behavior. It is reasonable to assume that the highest barrier will lead to the slowest dynamics, and this is described by the situation on the left panel of Fig. 5 (*AB* sequence). This is because the second barrier (counting from the deep intermediate state) is higher. For the case on the right panel in Fig. 5 (sequence *BA*), while the first transition is relatively slow it is not as slow as the second transition for the *AB* sequence. These arguments explain the effect of temporal order of mutations in cancer initiation.

**Figure 4.**
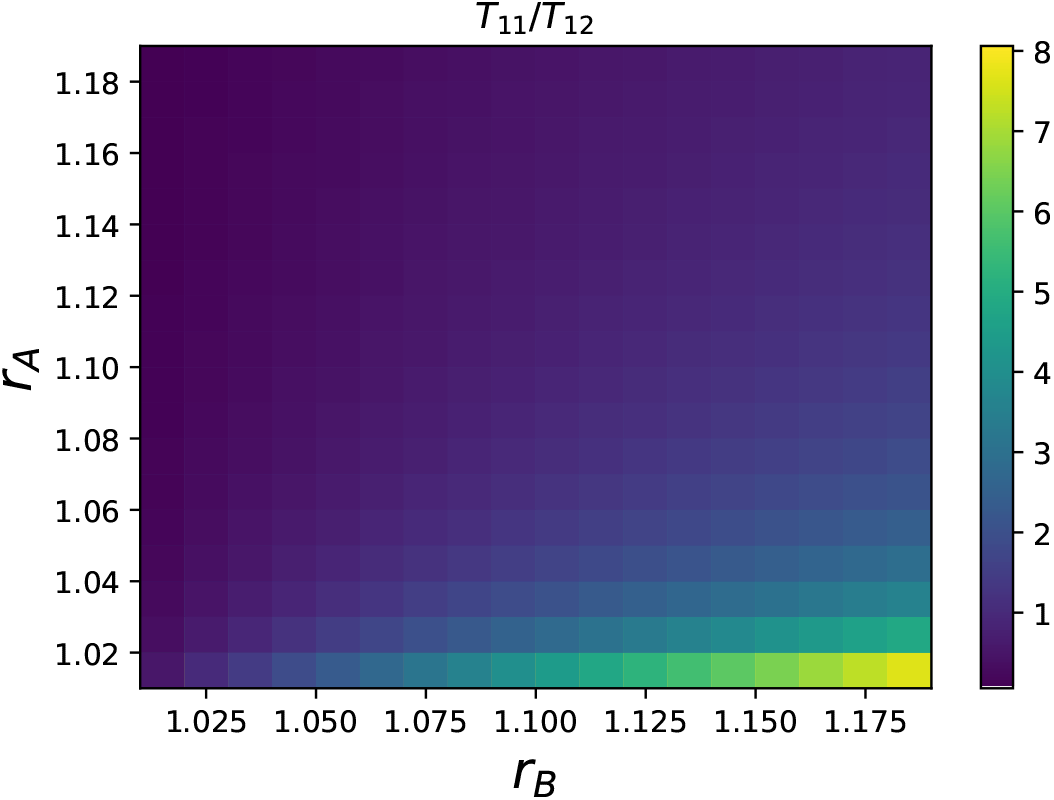
Heat maps for the ratio of the contributions to the overall fixation time *T*_11_/*T*_12_ as a function of different fitness parameters. In calculations, the *AB* mutation sequence and *N* = 1000 are chosen.

**Figure 5.**
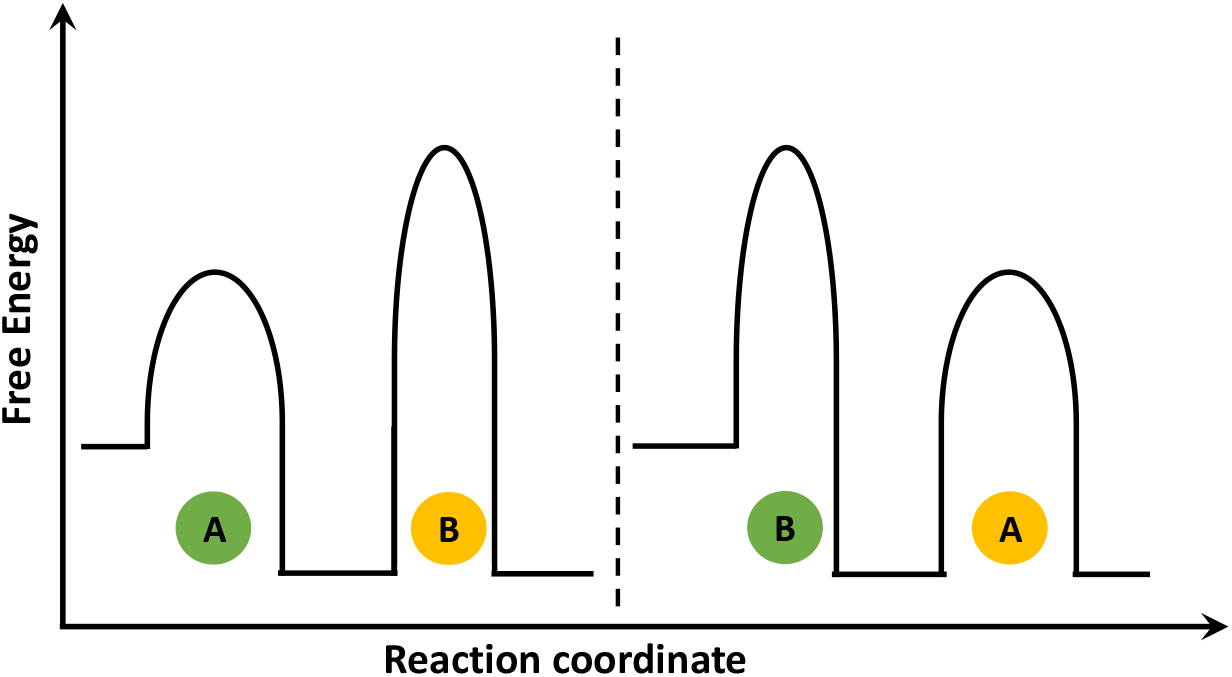
The effective “free-energy” landscapes for cancer initiation with two alternative sequences of mutations: left panel describes *AB* mutations (*r_A_* > *r_B_*), while right panel describes *BA* mutations (*r_B_* > *r_A_*).

In conclusion, we developed a theoretical model to quantitatively explain the effect of temporal order of mutations in cancer initiation. The appearance of tumor is associated with fixation of several mutations in the tissue compartment that originally had a fixed number of normal stem cells. Our idea is that the cancer initiation process can be viewed as a sequence of stochastic transitions between discrete states that are defined by different numbers of cells with one or two mutations. This allows us to evaluate properties of cancer initiation and compare different mutations sequences. Specifically, we analyzed in detail the cancer initiation dynamics after the fixation of two different sequential mutations. It is shown that the probability of cancer initiation is fully determined by the fixation of the first mutation. This also suggests that the probability of the tumor formation is higher if the first mutation is more advantageous than the second one. These theoretical results explain recent experimental observations that emphasize the special role of initial truncating mutations in the human cancers [13]. We also found that cancer initiation times depend on both mutations, and the fastest dynamics is observed if the second mutation is more advantageous. These theoretical predictions are able to explain striking observations that the reverse order of mutations delays the formation of tumor by many years or might even not lead to cancer at all [9]. One of the most surprising observations of our theoretical analysis is the apparent anti-correlation between the probability of cancer initiation and the speed of tumor formation, which contrasts with naive expectations. To better understand this result, we developed effective free-energy landscapes for cancer initiation dynamics that allowed us to clarify the mechanistic origin of this phenomenon. It should be noted that such anti-correlations are also frequently observed in other stochastic processes such as chemical reactions.

While the presented theoretical model provides a quantitative description of the effect of mutations order in the cancer initiation, it is crucial to discuss its limitations. It assumes that the second mutation does not start until the first one is fully fixed. This is known as a linear evolution in tumor formation [2]. However, there is a limited amount of experimental data that support the linear evolution in cancers [2,20]. Most studies favor a branching evolution picture in which different cellular clones with different mutations evolve in parallel in the growing tumors. At the same time, the linear evolution is a limiting case of the branched evolution when the mutation rate are very low, and this is frequently the case in many biological systems. In addition, there are arguments suggesting that early stages of cancer initiation can be well described by the linear evolution models [2]. Another simplification in our approach is the assumption that two mutations would lead to the formation of the tumor, while current data suggest that as many as 10 mutations are needed to drive some human cancers. Our method can be extended to multiple hits that follow the linear evolution. However, despite these limitations, the model still provides a clear physical picture for complex processes that are taking place during the formation of tumors.

## Acknowledgments

The work was supported by the Welch Foundation (C-1559), by the NSF (CHE-1664218), and by the Center for Theoretical Biological Physics sponsored by the NSF (PHY-1427654).

## Appendix

## I. Calculation of fixation probability and fixation times

One can define the corresponding first-passage probability *F_n_*(*t*) density functions to start from any state *n* and reach the site 2*N* at time *t*. The temporal evolution of these functions are governed by the following backward master equations,

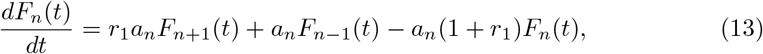

for 1 ≤ *n* < *N*, and

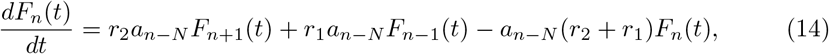

for *N* < *n* < 2*N*, while

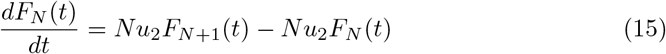

for *n* = *N*.

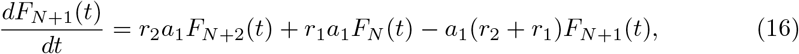

for *n* = *N* + 1. In addition we have the boundary condition *F*_2*N*_ = *δ*(*t*). It is convenient to solve this problem using the Laplace transformation, which changes the backward master equations:

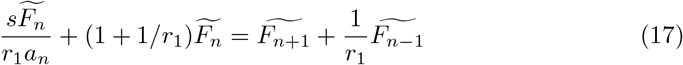

for 1 < *n* < *N*, and

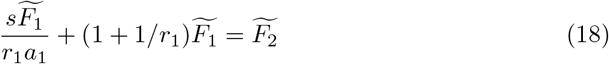

for *n* = 1.

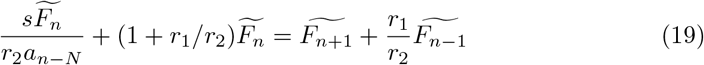

for *N* < n < 2*N*, and

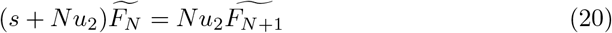

for *n* = *N*.

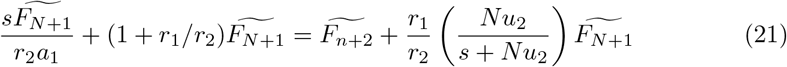

for *n* = *N* + 1. Because we are interested only in the fixation probabilities and fixation times, there is no need to obtain full analytical expressions for *F_n_*, but it is needed to determine the expansion of this function up to the linear term in *s*. Thus, we can write

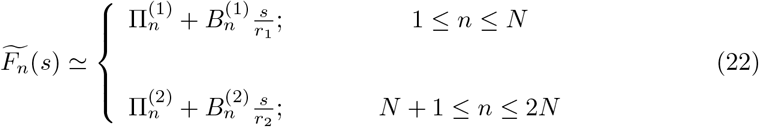

where 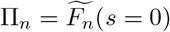, is the fixation probability starting from *n* single mutations, and the unknown parameters *B_n_* are related to the fixation times (viewed as conditional mean first-passage times) as

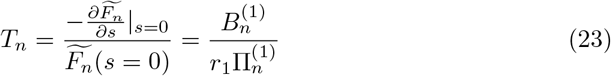

Note that because of boundary condition at *n* = 2*N* we have 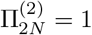 and 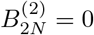. Substituting Eq. 22 into Eqs. 17 and 18 we obtain for the fixation probabilities

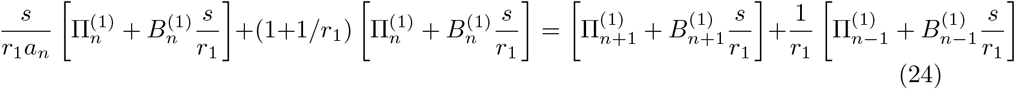

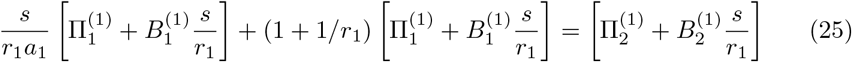

These expressions yield following equation for fixation probabilities:

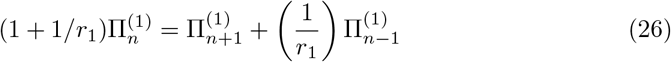

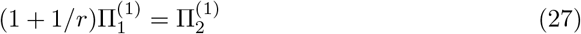

As shown in our previous work [23], these equations can be easily solved, leading to the following explicit expressions for the fixation probability,

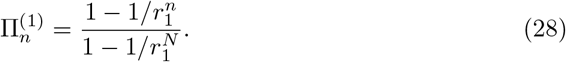

To proceed further we substitute Eq. 22 into Eqs. 19, 20, and 21

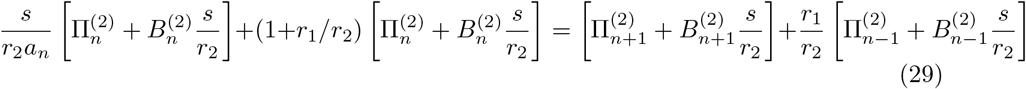

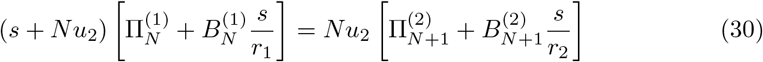

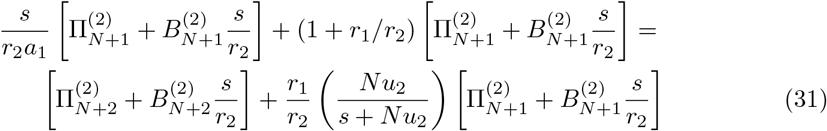

From these equations we obtain:

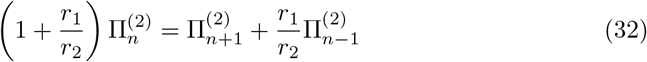

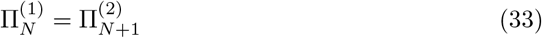

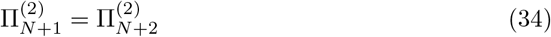

From eqn 28 and from boundary condition 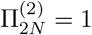 we obtain:

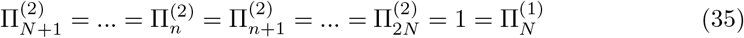

Thus fixation probabilities are given by:

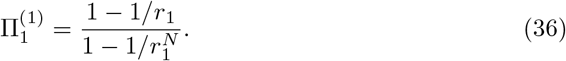

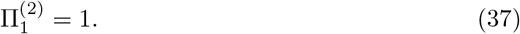

From Eqs. 24, the corresponding equations for parameters 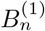 can be written as,

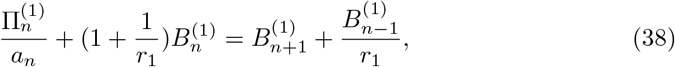

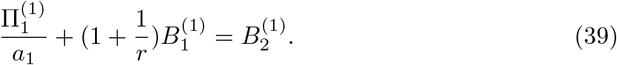

As was shown before, [23] 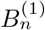 is given by:

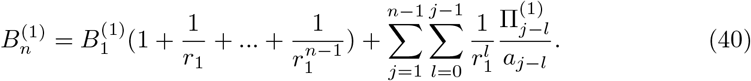

This expression is valid for any 1 ≤ *n* ≤ *N*. The constant 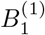 can be found from the boundary condition at *n* = *N*.

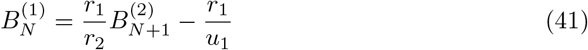

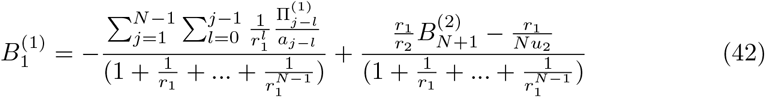

To proceed further we must calculate 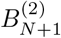. Similarly for 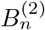 we obtain following recursion relation:

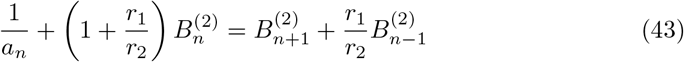

Now we define:

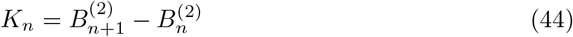

Thus Eqn. 43 takes the form:

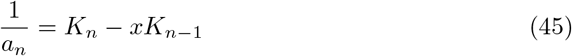

where 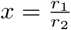. Using Eqn. 41, *K_N_* can be written as:

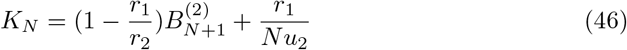

It can be shown that solution of recurrence relation 45 is given by:

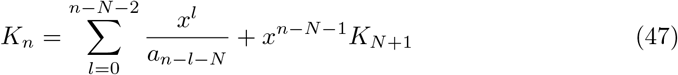

Thus 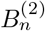 is given by:

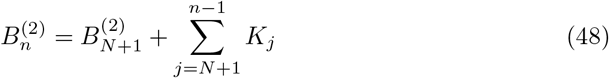

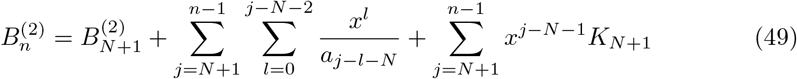

Now we can calculate *K*_*N*+1_:

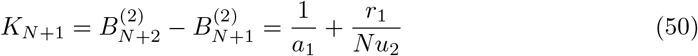

Combining Eqns. 50 and 49 yields:

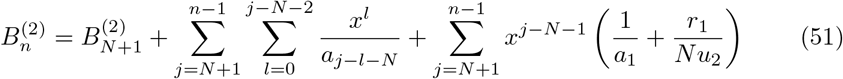

We can obtain 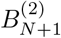 using boundary condition 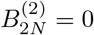:

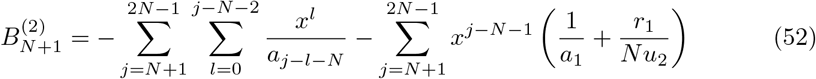

Substituting Eqn. 52 into Eqn. 42 yields:

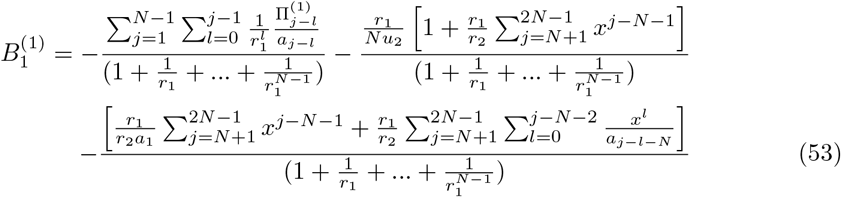

It can be shown that:

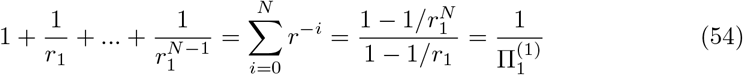

Now we can calculate fixation times using eqn. 23,

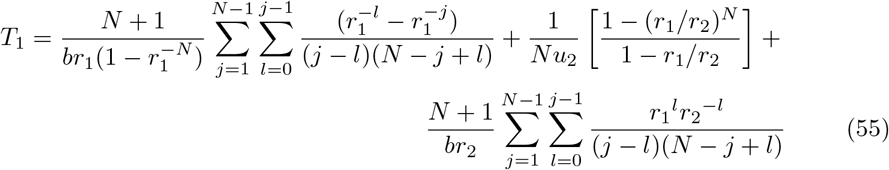

This expression can be further simplified by defining a new index *n* = *j* – *l*.

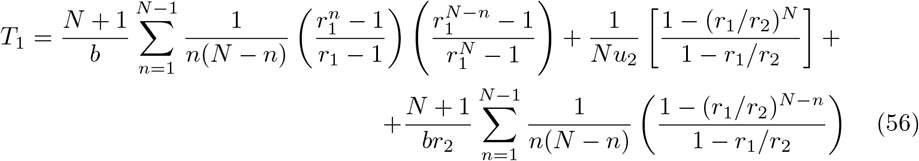

## II. Explicit expressions for fixation times for *N* → ∞

In general it is difficult to perform explicit summation for larger *N*. For *N* → ∞, we can convert summation to integration:

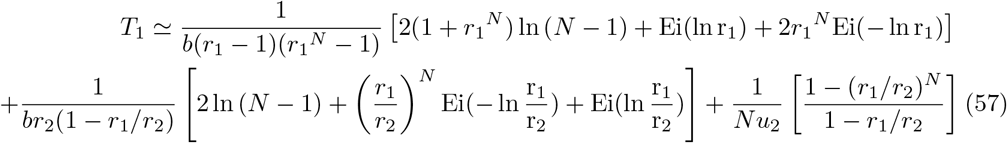

## III. Full exact solution of *F_n_* for *N* = 3

In this section we obtain a full exact solution of the first-passage probability density function *F_n_* for *N* = 3. This analytical solution is extremely useful, because it allows us to verify the general exact solution, which was obtained based on an steady state ansatz. As shown in figure 1, for *N* = 3 there are 6 states. Correspondingly we define the first-passage probability *F_n_*(*t*) functions to start from any state *n* and reach the site 2*N* at time *t*. The temporal evolution of these functions is governed by a set of backward master equations:

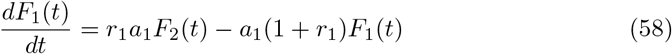

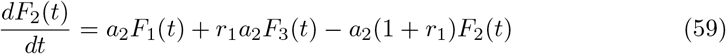

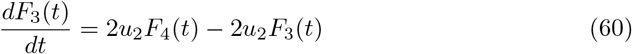

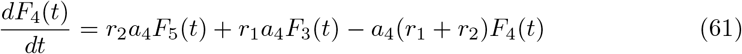

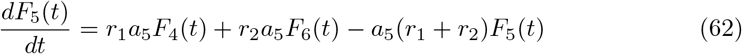

In addition we have the boundary condition *F*_6_(*t*) = *δ*(*t*). Performing Laplace transformation over these equations we get:

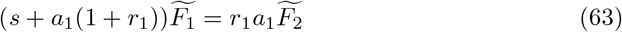

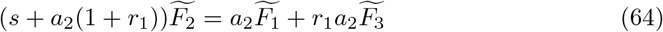

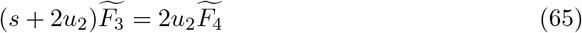

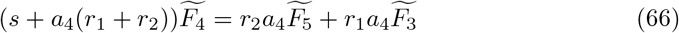

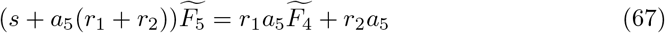

We note that because of symmetry we have 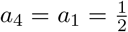 and 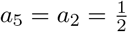. Solving this system of five equations and five unknowns yields 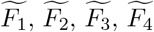 and 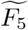. Specifically we need 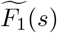 to calculate fixation time and fixation probability. After some algebra we obtain:

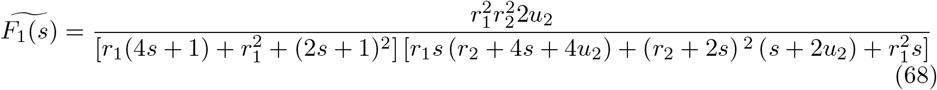

**Appendix II - Figure 1.**
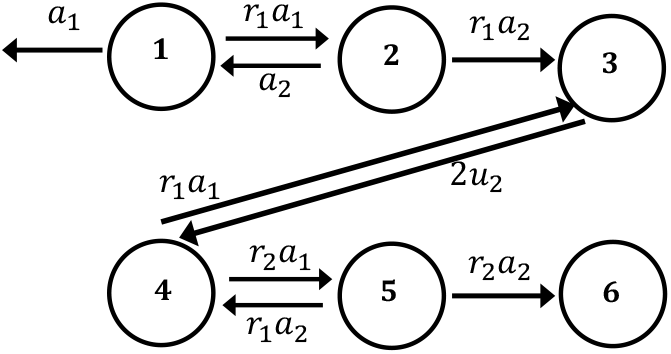
Discrete-state stochastic model of 2-hit mutation for *N* = 3.

Expanding this function in terms of *s* yields the fixation probability,

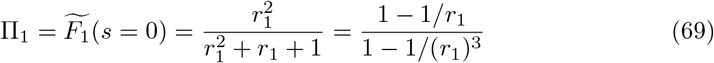

and, the fixation time:

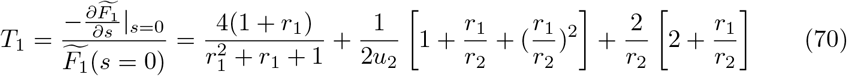

Similarly, the Taylor expansion of 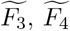 and 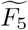 in terms of s yields the other fixation probabilities,

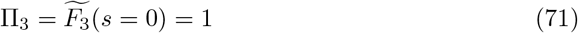

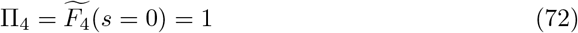

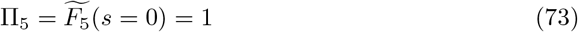

Now we expand 56 for *N* = 3 and after some algebra we obtain equation 70:

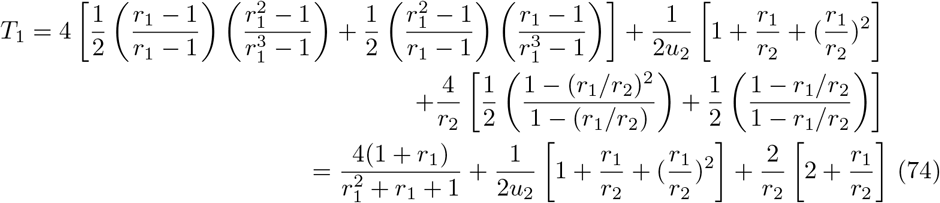

## IV. Effect of the number of stem cells

Another relevant parameter in the cancer initiation is the number of stem cells. Figure 2 shows how varying *N* influences the differences in the cancer initiation probabilities and times. The differences are relatively small for low number of cells, but they start to increase and eventually saturate at *N* ≫ 1. The differences are stronger for the fitness parameters that are closer to unity. This can be easily explained for the fixation probabilities because Π_1_ = 1 – 1/*r*_1_ in the limit of *N* → ∞. In this limit, from Eq. (57) the fixation time can be approximated as *T*_1_ ~ ln *N* with the coefficient that depends only the fitness parameters *r*_1_ and *r*_2_, and this leads to the constant ratio of the fixation times for different mutation sequences.

**Appendix IV - Figure 2.**
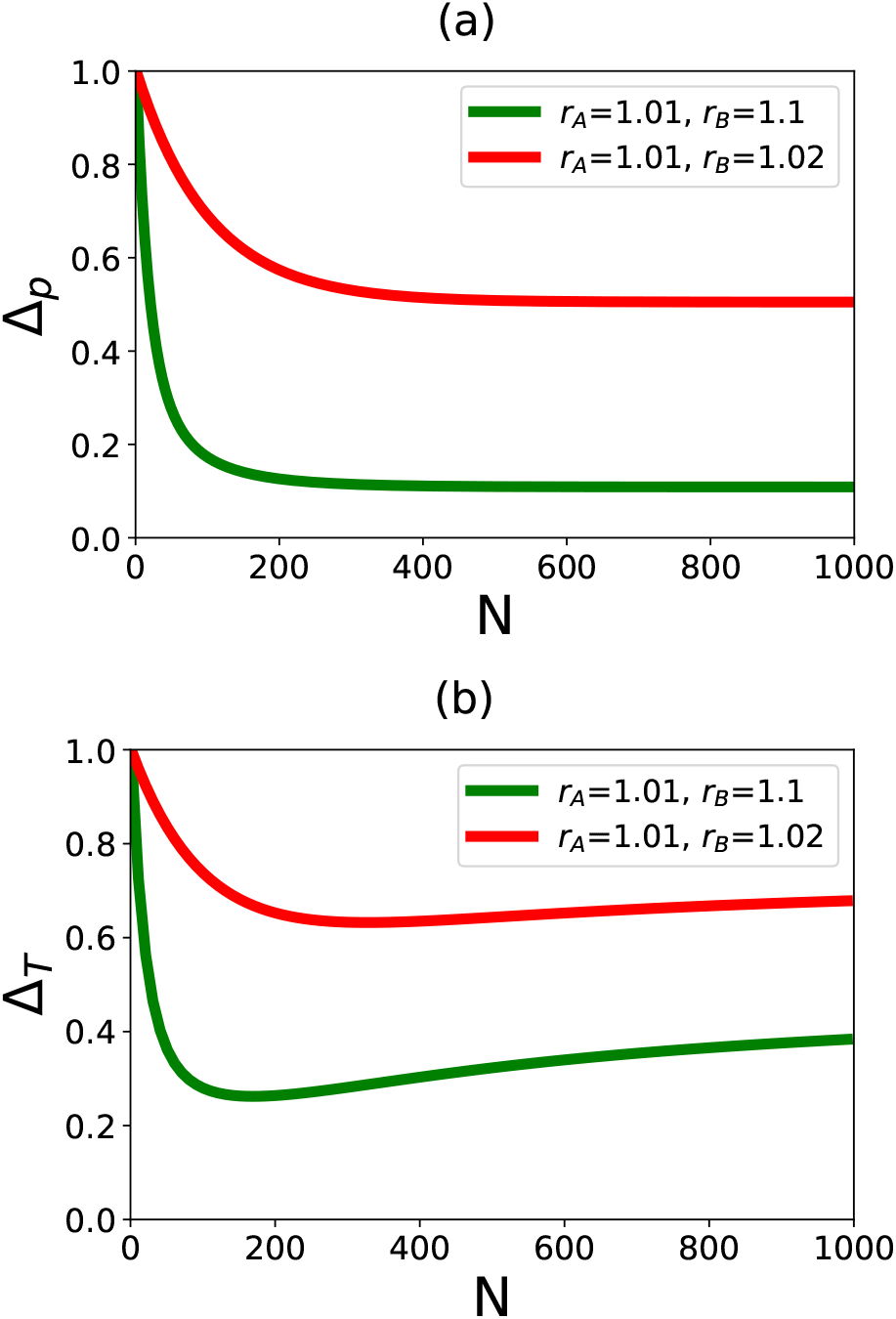
(a) The ratio of fixation probabilities Π_*AB*_/Π_*BA*_ as a function of the number of cells *N*. (b) The ratio of fixation times *T_AB_/T_BA_* as a function of the number of cells *N*. For calculations, *u*_2_ = 10^-4^ is used.

